# Plants species’ influence on rhizosphere microbial communities depends on N availability

**DOI:** 10.1101/2021.11.23.469737

**Authors:** Teal S. Potter, Brian L. Anacker, Amber C. Churchill, William D. Bowman

**Affiliations:** Ecology and Evolutionary Biology, University of Colorado, Boulder, CO 80309; Department of Crop and Soil Sciences, Washington State University, Pullman, WA 99164, USA; Boulder County Open Space, Boulder, CO, 80301; Department of Ecology, Evolutionary Biology and Behavior, University of Minnesota, St. Paul, MN 55108; Institute of Arctic and Alpine Research, Boulder, CO 80309

**Keywords:** plant-microbe interactions, *Poa*, rhizosphere, plant effects, nitrogen deposition

## Abstract

**Purpose:** Plants and soil microbes both influence how ecosystems respond to environmental change. Yet, we lack the ability to generalize how plants and soil microbes influence each other in the same or varying soil conditions. This limitation thwarts ecologists’ ability to understand and predict effects of environmental changes such and elevated anthropogenic nitrogen (N) deposition. Accordingly, we examined the specificity of plant species’ influence on soil microbial community composition.

**Methods:** We tested (1) whether congeneric grass species have unique effects on soil microbial communities, (2) how relative abundances of microbial taxa can be explained by *Poa* phylogeny, plant traits, and range-wide traits (annual temperature and soil pH), and (3) whether N addition alters associations between *Poa* species and soil microbes, and (4) whether the magnitude of microbial community change in response to elevated N can be explained by plant growth responses to N. We conducted a greenhouse experiment with seven *Poa* species and native soils.

**Results:** We found that individual *Poa* species were associated with different soil fungi but similar soil bacteria. Differences in microbial composition were not attributable to *Poa* phylogeny, plant traits, or range-wide traits. Nitrogen addition enhanced the unique effects of *Poa* species on fungal and bacterial community compositions.

**Conclusion:** These results demonstrate how ecological interactions of related plant species vary depending on resource supply, revealing important context dependency for accurately predicting microbially-mediated nutrient cycling and ecosystem responses to changes in nutrient availability.

## Introduction

The interaction between plant roots and soil microbes affects the fitness of individual plants, plant community composition, and important ecosystem services, such as nitrogen (N) cycling and carbon (C) storage. Plants and soil microbes reciprocally influence each other by providing or withholding resources (Kiers et al. 2011). These rhizosphere interactions are often mediated by plant root exudates that impact microbial colonization, microbial signaling, rhizosphere acidity, and nutrient uptake (Nguyen 2003; Bais et al. 2006; Hartmann et al. 2009). To date our understanding of rhizosphere interactions indicates a high degree of complexity, where plant species differ in which soil microbes they associate with, and that these differences are in turn dependent on environmental factors like temperature, soil moisture, soil pH, and which microbial taxa are nearby. Finding ways to generalize plant-soil interactions across plant species or soil types remains a persistent challenge. A particular need is to learn how changes in nutrient availability, such as with anthropogenic nitrogen (N) deposition, alters plant-microbe interactions differently among plant species (Classen 2015).

Among plant species, variation exists in the types and amounts of root exudates they produce may lead to specificity in plant-soil microbe associations (Bardgett et al. 1999; Berg and Smalla 2009; Reese et al. 2018). For example, when different plant species are grown in a common soil, species-specific root trait variation can facilitate the assembly of different microbial communities from the same pool of soil microbes (Haichar et al. 2008; Philippot et al. 2013). Microbial community assembly and activity are also influenced by physical soil properties (Girvan et al. 2003; Singh et al. 2007; Lauber et al. 2008). Moreover, when plants alter root-associated microbial community composition in a way that promotes or inhibits the growth of conspecific plants, a positive or negative feedback is initiated (Bever 1994). The direction and magnitude of feedbacks are largely attributed to changes in the structure and function of microbial communities (Callaway et al. 2004; Chapman et al. 2006; Bever et al. 2010). However, few experiments have attempted to systematically characterize differences in plant species’ effects on microbial communities (Grayston et al. 1998; Pennings et al. 2005; Appuhn and Joergensen 2006; Veresoglou and Rillig 2014), which could aid in uncovering drivers of plant-soil feedbacks.

Using plant traits, such as plant height and specific leaf area, as potential indicators of *belowground* biology is an area of active investigation (Veresoglou and Rillig 2014; Mehrabi and Tuck 2015; Münzbergová and Šurinová 2015; Leff et al. 2018). While plant traits and phylogeny are sometimes proxies for unmeasured traits often explain variation in *aboveground* ecological interactions (Gilbert and Parker 2010; Burns and Strauss 2011), plant traits and phylogeny do not appear to exhibit consistent relationships with belowground soil biota or soil chemistry (Burns and Strauss 2011; Anacker et al. 2014; Cantarel et al. 2014; Moreau et al. 2015; Fitzpatrick et al. 2017). Selecting the appropriate taxonomic scale may be essential for detecting variation in plant traits that correspond to plant effects on soil. For example, at the taxonomic scale of different genotypes within a plant species (i.e., intraspecific), plant trait variation may often be too small to detect a correlation with belowground microbial communities (Wagner et al. 2016; Emmett et al. 2017; Leff et al. 2017). On the other extreme, at the taxonomic scale of different species (from different families or different orders) variation in plant traits may be too large to link traits to belowground interactions (Bardgett et al. 1999; Smalla et al. 2001; Innes et al. 2004; Garcia et al. 2005; Anacker and Strauss 2016). In contrast, the understudied, intermediate taxonomic scale of *genus* may be “just right” for detecting trait differences that matter to plant-soil microbe interactions, but, of course, this general principle may not hold for all genera.

The context-dependency of plant-soil interactions may have important consequences for nutrient cycling and ecosystem responses to environmental change (Shade et al. 2012; Classen et al. 2015). For example, anthropogenic N deposition (Vitousek et al. 1997; Galloway et al. 2008) influences plant species’ abundances and diversity as well as soil biota and chemistry in many affected ecosystems (Galloway et al. 2008; Bobbink et al. 2010; Simkin et al. 2016; Payne et al. 2017). The separate effects of elevated N on plants (Tilman and Wedin 1991; Vitousek and Howarth 1991; Stevens et al. 2004) and soil microbes (Treseder 2004; Ramirez et al. 2010; Geisseler and Scow 2014) have been studied extensively. However, rarely are the effects of N on plants and soil microbes studied together (Bardgett et al. 1999; Moreau et al. 2015).

Plant species may differ in their response to increased N (Stevens et al. 2004; Pennings et al. 2005) in part due to changes in resource allocation in root tissue and concurrent changes in soil microbial populations. For example, plant species with strong growth responses to added N may also increase allocation of carbon (C) to roots (Rinnan et al. 2007), which in turn alters microbial community composition (Paterson et al. 2007). An increase in soil N can also directly affect microbial community composition, disrupting or obscuring plant species’ effects on microbial community composition (Kennedy et al. 2004). Investigating how the dynamics of plant-microbe interactions are mediated by an environmental change, such as N deposition, will help disentangle the direct and indirect effects of plants and N on soil microbes.

We conducted a greenhouse experiment to examine plant-microbe interactions with and without simulated N deposition using seven grass species in the genus *Poa*. We tested four hypotheses about plant-microbe interactions: (1) Congeneric plant species have unique, non-random effects on rhizosphere microbial composition, (2) Differences in microbial community composition among plant species can be explained by plant traits and phylogeny as a proxy for unmeasured traits, (3) N addition alters the distinct effects of plant species on microbial community composition, and (4) the magnitude of microbial community change in response to elevated N can be explained by plant growth responses to N.

## Methods

To assess whether closely related grass species have unique effects on rhizosphere microbial composition, seven grass species in the genus *Poa* were used in a greenhouse experiment. By growing plants of different species in a uniform field soil we ensured that each pot received a similar starting composition of soil microbes to be able to statistically distinguish unique effects of *Poa* species. Using a no-plant treatment (pots with soil but no plants) and a N addition treatment for all plant species and no-plant controls enabled distinguishing among N effects and plant effects on soil microbial community composition.

We identified a set of plant traits to test alternative hypotheses associated with the role of plant traits that in explaining plant species’ effects on microbial communities. We used plant height (longest leaf), dry shoot mass, dry root mass, and specific leaf area (SLA) from ambient N treatments to serve as species’ trait measurements. We chose these plant traits as potential indicators of plant species’ effects on soil microbial communities because they are relevant for plant nutrient use and allocation (Aerts and Chapin 1999; Grassein et al. 2015). These traits distinguish ecological and evolutionary variation among plant species, which sometimes capture trade-offs in growth strategies (e.g. SLA)(Funk et al. 2017). Additionally, SLA and height differ between native and invasive plant species (Van Kleunen et al. 2010) which are both represented in this study. It is not yet known which—if any—plant traits can explain differences in plant-N-soil microbial relationships. Thus, we used a suite of common traits that generally describe different aspects of ecological niches (Funk et al. 2017). Our intent was not to perform an exhaustive analysis of potential traits to find the best surrogate for exudate composition that drives variation in microbial communities among plant species. Instead, our goal was to test alternative hypotheses (other than phylogenetic relatedness). Additionally, we also chose to test two species-level traits that represent environmental variation among the *Poa* species’ geographical ranges to determine if plant effects on soil microbes may be explained by plant species’ adaptations to particular environments. We obtained geographical and elevation ranges for each plant species from the GBIF database (GBIF 2017) and used this information to estimate species’ means for soil pH in their home ranges from the ISRIC Soil Data Hub (ISRIC - WDC Soils 2017) and mean annual temperature in their home ranges from PRISM (Climate Group 2017).

The seven species used in the experiment were *Poa alpina, P. arctica, P. compressa, P. glauca, P. nemoralis, P. pratensis*, and *P. reflexa*. This set of plant species was chosen due to a fully resolved phylogeny (Gillespie et al. 2007), relatively low rates of hybridization, and variation in phylogenetic distance among the seven species. Choosing a large genus was also convenient for locating species in a small geographic area. Additionally, a subset of these species has similar plant traits, habitats, or both, which allowed for determining whether these factors correlate with plant effects on microbial community composition. The three alpine species (*P. alpina, P. arctica*, and *P. glauca)* are distantly related within this set of seven species but coexist in the same plant community, providing an opportunity to determine whether phylogenetic signal is a better or worse indicator of microbial composition than plant traits or environmental niches. Two species, *P. pratensis* and *P. compressa*, are non-native, widespread, and may possess similar traits for successful invasion in the Colorado Front Range. Like the alpine species, these species coexist but are not closely related in the pruned phylogeny. All seven selected *Poa* species are perennial and rhizomatous.

*Poa* species were identified using the two leading taxonomic guides for the region (Weber and Wittmann 2012; Ackerfield 2015) during the summer of 2014. Populations of the *Poa* species used in the study are in Boulder County, CO, USA and spanned an elevational gradient from low montane (elevation 1764 m, lat 40.124012° N, long -105.30666° W) to alpine (elevation 3466 m, lat 40.052486° N, long -105.582467° W). Seeds were collected from approximately 20 inflorescences presumed to include several genets. These inflorescences were distributed across the geographic are of source population of each species. Voucher specimens are archived at the University of Colorado’s herbarium under the following accession numbers: *P. compressa* 02386647, *P. nemoralis ssp. Interior* 02388189, *Poa reflexa* 02388262, *P. alpina* 02388288, *P. arctica* 02388197, and *P. glauca* 02388270.

Nine liters of combined bulk and loosely root-attached rhizosphere soil was collected from the collection sites for each *Poa* species’ population in September. Soils from the seven *Poa* populations were stored between 10-21°C in open one-gallon plastic bags for less than 1 week before soil was mixed separately for each soil type. Soils were mixed for five minutes, sieved (2 mm), and then an equal volume of soil from each site was combined and thoroughly mixed for five minutes to create a uniform soil type. This uniform soil was created so that each pot would have a similar starting composition of soil microbes, including microbes found from each of the seven *Poa* species’ ranges. Conical pots (164 ml) were filled with the mixed soil to within 2 cm of the top of the pots.

Seeds from each species were surface sterilized with 10% bleach, and seeds were planted in half of the prepared pots. The number of seeds planted per pot (<10) was determined by a germination trial in petri dishes. Following germination, plants were thinned to a single seedling per pot. Tools were sterilized with bleach before thinning a new pot to minimize contamination of microbes between pots. Where no plants germinated in a pot (approximately 5% of pots) an extra seedling from another pot was surface sterilized in 7% bleach for 10 minutes, rinsed with DI water and transplanted.

Four treatment levels were established. These treatments were soil only (hereafter, no-plant controls), soil with N addition, soil with one plant and no N addition, and soil with one plant and N addition. These treatments allow for distinguishing plant effects, N effects, and the interactive effects of plants and N on soil microbes. We used ten replicates per treatment per plant species for a total of 160 pots.

A 595 watt LED panel was installed above the racks to supplement low levels of natural light (augmented light produced 360 µmol/photons/m^2^ below light to 215 µmol/photons/m^2^ on edge of racks on a cloudy afternoon). Racks were randomly rotated under the LED panel every week to minimize the effects of light intensity variation. Lights were on for 13 hours a day, which was increased to 14 hours a day halfway through the experiment to mimic natural photoperiod of the growing season. Greenhouse temperatures ranged from 10-21°C, which is well within the range all *Poa* species experience in the field during their growing season.

Four weeks after germination, half of the planted pots and half of the soil-only pots began receiving N treatments twice each week as a solution of 1mmol NH4NO3 for a total of 11 N applications. The young plants were watered with 20 ml of tap water (with or without N), which was poured onto each pot’s soil twice per week to maintain constant soil moisture throughout the experiment (Fig. S6). The N application rate was equivalent to 70 kg N ha^−1^. Tap water contained <0.05 mg N L^-1^ in the form of nitrate.

After 17 weeks—an intermediate growing season length between the lowest and highest elevations of *Poa* populations—the plants were harvested. This timing also allowed for harvesting plants before they became pot bound. Plant height (length of longest leaf) was measured on live plants as a proxy for plant growth before plants were harvested. For each pot, bulk soil was collected for soil pH measurements. Soil samples were also collected from all pots for microbial community analysis. From no-plant control pots, a sample was pooled from multiple parts of the pot using bleach-sterilized metal spatula. Rhizosphere soil from pots with plants was collected by shaking soil from roots, and then the remaining soil adhering to roots was then shaken and gently massaged from roots onto a clean sheet of weigh paper. This rhizosphere soil was then transferred to a sterile tube and kept on ice until samples could be moved to a -20°C freezer at the end of each harvest day. Roots were then washed, and roots and shoots were separated and placed in labeled envelopes and dried at 60°C for 48 hours to assess dry mass. To measure SLA, one representative leaf per plant was attached to a white piece of printer paper using clear tape, which was scanned the same day and used for leaf area measurement using the software program ImageJ (Rasband W. S., ImageJ, US National Institute of Health, Bethesda, MA). Because individual plants are small, leaves used for leaf area were then removed from the paper, dried and added to individual plant’s dry shoot mass. Soil pH was measured using a 2:1 soil:tap water slurry which was shaken for 30 minutes and measured using a Beckman 340 pH probe (ICP-AES; Thermo Electron, Waltham, MA, USA).

DNA from soil samples was extracted using MoBio Laboratories PowerSoil DNA isolation kit (Mo Bio Laboratories, Inc.), following the manufacturer’s instructions. To ensure that a representative sample of microbial taxa from each plant’s rhizosphere was used for DNA extraction, an additional step was taken in which 0.25 g from each soil sample was combined with 0.4 ml ultra-pure water in a sterile microcentrifuge tube and shaken before subsampling. From the slurry, 0.25 g were transferred to a well in 96 well power bead plates to begin the normal DNA extraction protocol.

To obtain microbial composition for each no-plant control pot or *Poa* rhizosphere, the V4 region of the16S ribosomal RNA gene and the ribosomal internal spacer ITS1 were amplified using unique barcodes for each sample for multiplexing with 16S and ITS primers to amplify bacterial and fungal DNA respectively. PCR product was pooled after performing duplicate reactions and cleaned using a SequalPrep Normalization kit (Thermo Fischer Scientific Inc.). The cleaned and pooled amplicon was then sequenced on an IlluminaMiSeq running 2 × 250 bp sequences. Sequences were demultiplexed, and quality filtering and phylotype clustering were conducted using the UPARSE pipeline (Edgar 2013). Raw sequences were then mapped to phylotypes (97% similarity) after removing singletons. The GreenGenes database was used to obtain taxonomy classifications for OTUs using the RDP classifier. Datasets were then checked for contamination, and samples were rarefied to 3997 sequences per sample in the 16S dataset and 1000 sequences per sample in the ITS dataset to account for differences in sequencing depth.

To estimate pair-wise phylogenetic distances among the seven *Poa* species we created a molecular phylogeny for the seven plants species based on four genes: ITS, rbcL, matK, and trnH-psbA. Branch length calibration was conducted in BEAST using the topology from Gillespie et al. 2007 a constraint and calibration points for the root node of 50 Ma and an arbitrary standard deviation of 1.0 Ma with a normal distribution. We ran an MCMC chain for 10 million generations, sampling every 1000 generations. Convergence of the posterior distribution was checked using TRACER v1.5. From the combined BEAST posterior distribution of 10,000 trees, we removed the first 2,000 trees as burnin and combined the remaining 8,000 trees into a single maximum credibility tree using TreeAnnotator v1.8 (Fig. S1). Phylogenetic distance among species pairs were extracted from a phylogenetic distance matrix calculated using the “cophenetic” function in the Picante library.

### Data Analyses

Statistical analyses were performed in R (R Core Team 2016). The package ‘mctoolsr’ (http://leffj.github.io/mctoolsr/) was used to aid in formatting of multivariate datasets.

To determine which traits if any explained observed variation in microbial community composition, we used linear regressions to test for relationships between microbial community composition and plant traits, phylogenetic distance, and environmental predictors separately. For regression analyses, microbial community composition was necessarily transformed from a community matrix of relative abundances of taxa to univariate data by calculating Bray Curtis dissimilarities. This dissimilarity matrix was then collapsed into a three-column data frame with two identifier columns (e.g. *P. artica* 1 with N, *P. arctica* 1 without N) associated with a single value from the dissimilarity matrix. Dissimilarities were then regressed against between *Poa* species differences in trait values.

A similar procedure was used to prepare data for linear regression to test hypothesis 3, whether plant species’ growth responses to add N impact the degree of microbial community change. For this test, we calculated plant species’ growth response to N by subtracting the mean plant mass among technical replicates within a *Poa* species from each of the 10 technical replicates of the same *Poa* species that received N addition. This resulted in 10 values per species to include in the analysis. Then, to calculate microbial community responses to N within *Poa* species we constructed a Bray Curtis dissimilarity matrix with only relevant contrasts (within *Poa* species ambient N means and N addition technical replicates)(Fig. S2).

We used PerMANOVAs using the adonis function in the vegan R package to test hypotheses one and three; to determine whether variation in bacterial and fungal community composition could be explained by plant species and N addition respectively. Differences in microbial community composition were calculated by comparing Bray-Curtis distances of the relative abundance of all OTUs per treatment. Microbial communities associated with no-plant controls were included in a model to first determine if there was a general effect of all plants on soil communities regardless of plant species. Then no-plant controls were removed, and the analysis was run again to determine whether microbial communities differed among *Poa* species. PerMANOVA was also used to test how N addition altered plant effects on rhizosphere microbial communities by determining differences in microbial communities among plant species, between N treatments and the potential interaction between plant species and N addition.

After determining that pH differed among soil samples at the end of the experiment, we recognized the potential for pH to have influenced microbial community composition differently among samples. We thus ran a variance partitioning model with N treatment and soil pH as potential mediators of microbial community composition. We attributed variation in microbial community composition to either N, pH, or both, while including plant species information using the varpart function in the vegan package. Using this method, explanatory variables were run as partial regressions with community dissimilarity matrices.

Linear regression was used to test hypothesis four: the magnitude of microbial community change in response to elevated N can be explained by plant growth responses to N. Data preparation is described above. An interaction term (plant height N response * *Poa* species) was included to examine the potential *Poa* species to have different relationships with microbial community responses.

## Results

We first tested whether the plant traits we measured were correlated with phylogenetic relationships among *Poa* species before testing plant species effects on microbial composition. We found no relationship between phylogenetic distance and trait similarity for root mass (Table 1). However, there was a positive linear relationship between phylogenetic distance and plant height, which was measured as the longest leaf on the plant (Table 1).

**Table 1.**
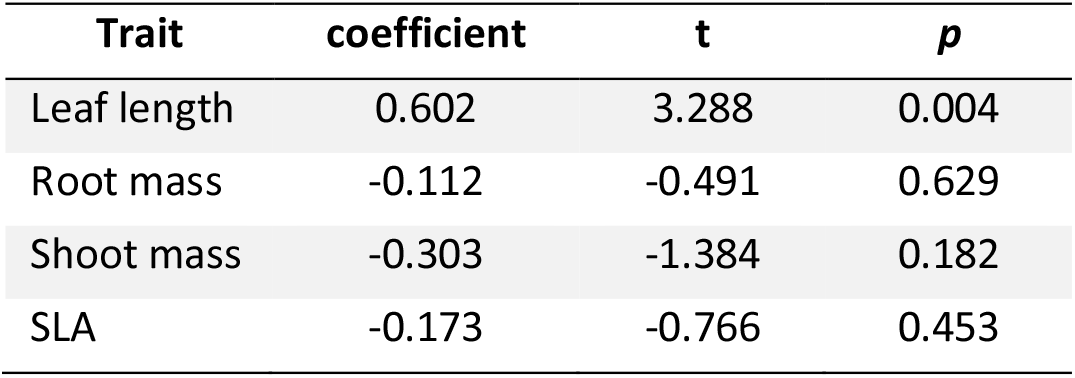
Pearson’s correlation test results of plant measurements among *Poa* species versus phylogenetic distance distances among species

### Plant species’ effects on microbial composition

*Poa* species and no-plant controls had unique effects on fungal community composition but not bacterial community composition (Table 2). When no-plant controls were removed from the analysis to test for a plant species effect, *Poa* species again had unique effects on fungal community composition but not bacterial community composition (Fig. 2, Table 2). Consequently, only fungal data were used to determine whether hypothesized predictors could explain differences in microbial community structure among *Poa* species.

**Table 2.**
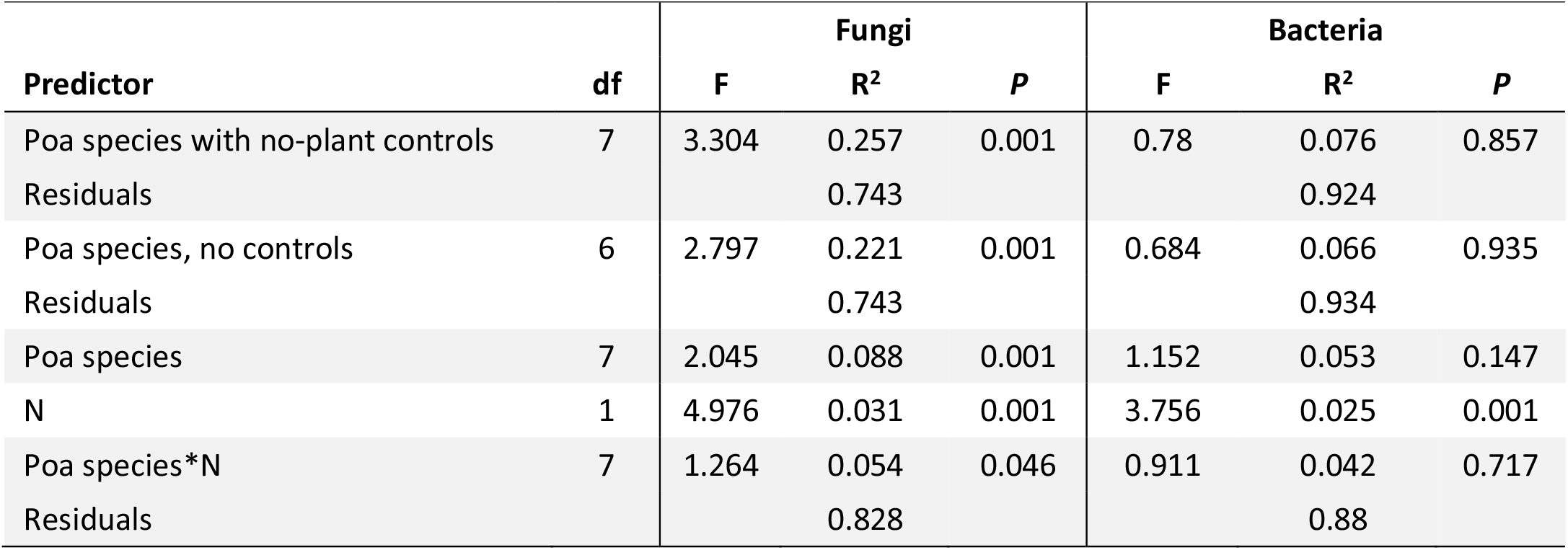
perMANOVA results for hypotheses tests 1 and 3

| Predictor | df | Fungi |  |  | Bacteria |  |  |
| --- | --- | --- | --- | --- | --- | --- | --- |
|  |  | F | R<sup>2</sup> | P | F | R<sup>2</sup> | P |
| Poa species with no-plant controls | 7 | 3.304 | 0.257 | 0.001 | 0.78 | 0.076 | 0.857 |
| Residuals |  |  | 0.743 |  |  | 0.924 |  |
| Poa species, no controls | 6 | 2.797 | 0.221 | 0.001 | 0.684 | 0.066 | 0.935 |
| Residuals |  |  | 0.743 |  |  | 0.934 |  |
| Poa species | 7 | 2.045 | 0.088 | 0.001 | 1.152 | 0.053 | 0.147 |
| N | 1 | 4.976 | 0.031 | 0.001 | 3.756 | 0.025 | 0.001 |
| Poa species*N | 7 | 1.264 | 0.054 | 0.046 | 0.911 | 0.042 | 0.717 |
| Residuals |  |  | 0.828 |  |  | 0.88 |  |

Statistically significant changes in relative abundances of 8% of the 129 tested common fungal families (Table S2) occurred with at least one *Poa* species relative to no-plant control communities after applying Bonferroni alpha corrections for multiple tests. Changes in abundances of these taxa likely contributed to the observed change in overall community composition.

Neither plant traits (plant height, root mass, shoot mass, and SLA) nor phylogenetic relatedness explained differences in *Poa* species’ effects on fungal community composition (Table 3). Phylogenetic distance showed linear relationships with only three of the 171 common fungal families tested after applying Bonferroni alpha corrections for multiple tests (Table S3). Additionally, neither of the two range-wide traits we tested, soil pH and mean annual temperature, exhibited a linear relationship with fungal community composition (Table 3).

**Table 3.** Linear regression results indicate that neither *Poa* phylogenetic distance nor *Poa* traits predict fungal taxonomic composition

| **Trait** | **coefficient** | **SE** | **R<sup>2</sup>** | **P** |
| --- | --- | --- | --- | --- |
| Phylogenetic distance | -0.021 | 0.022 | 0.003 | 0.344 |
| Height | 0.029 | 0.031 | -0.008 | 0.371 |
| Root mass | 0.03 | 0.038 | -0.02 | 0.444 |
| Shoot mass | -0.052 | 0.033 | 0.069 | 0.132 |
| SLA | -0.041 | 0.027 | 0.063 | 0.142 |
| MAT in species' ranges | 0 | 0 | -0.052 | 0.929 |
| Soil pH in species' ranges | -0.002 | 0.003 | -0.028 | 0.509 |

### Nitrogen addition influences Poa species effects on soil microbial communities

While individual *Poa* species did not show unique effects on bacteria composition in ambient N conditions, *Poa* species did show a consistent effect on bacterial community composition that was dependent on N treatment (Table 2, Fig. 1D). The same was true for fungal community composition, as indicated by a significant interactive effect of N addition and *Poa* species (Table 2, Fig. 1C). Differences in bacterial and fungal community composition remained significantly different among *Poa* species and the two N treatments after microbial data from no-plant control pots were removed from the analysis (Table 2). Differences in community composition between N addition and ambient N treatments can be attributed to increases in the relative abundances of a subset of microbial populations that showed preference for either ambient N or elevated N conditions (Figures 2, S3, S4).

**Fig. 1.**
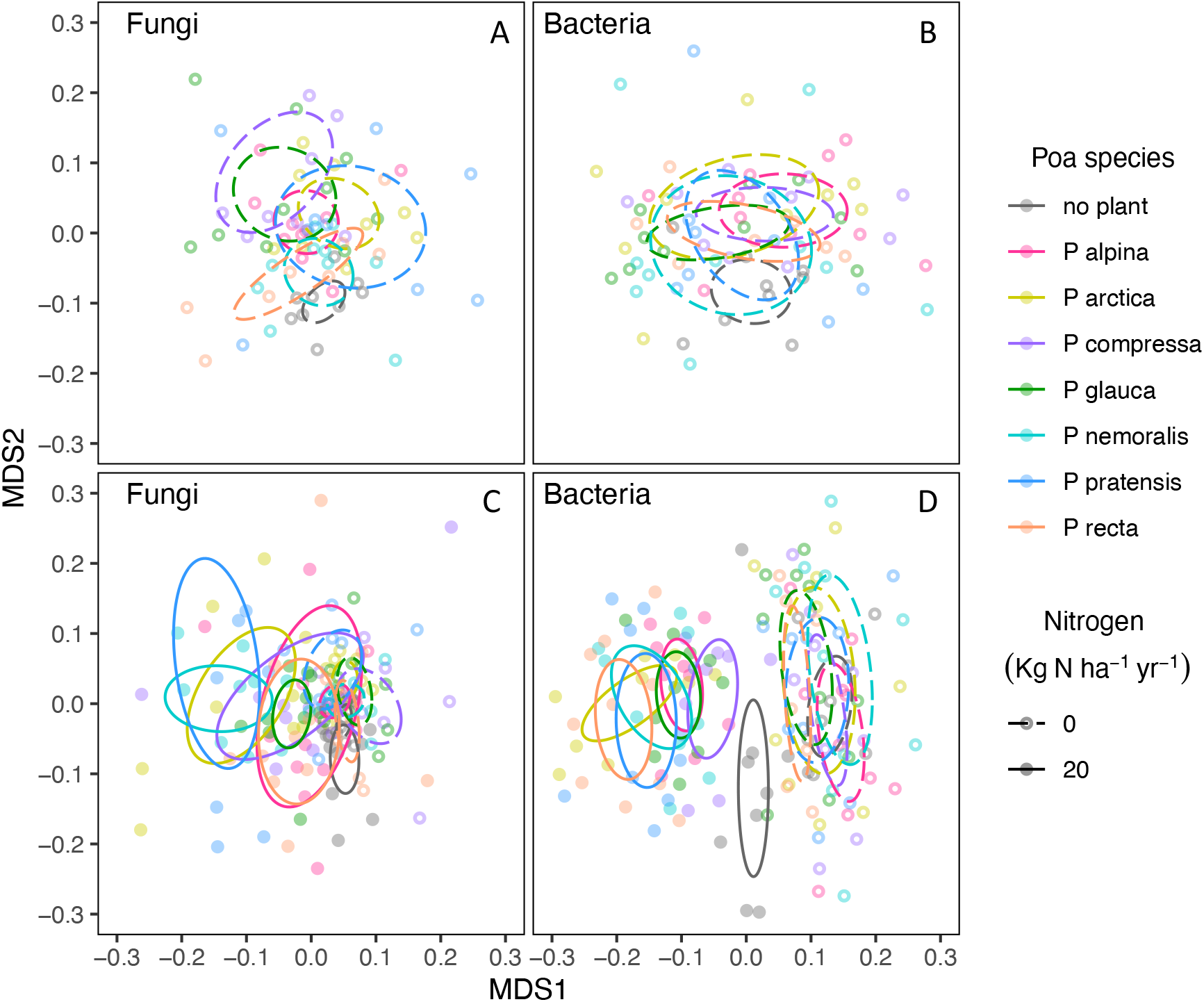
NMDS ordination showing microbial community compositional differences among rhizosphere samples and no-plant control samples for all *Poa* species. Orbitals are 95% confidence limits for each *Poa* species, where dashed lines represent non-fertilized and solid lines represent N fertilized communities. Panel A) shows fungal composition from non-fertilized pots (stress = 0.249) and B) shows bacterial composition in non-fertilized soils (stress = 0.208), while shows C) fungal composition from both N fertilized and non-fertilized pots (orbitals mostly overlap each other)(stress = 0.237) and d) shows bacterial composition from both N fertilized and non-fertilized pots (stress = 0.195). Rhizoshpere microbial community composition differs significantly among *Poa* species in panels A, C, and D (perMANOVAs, *P* < 0.05).

**Fig. 2.**
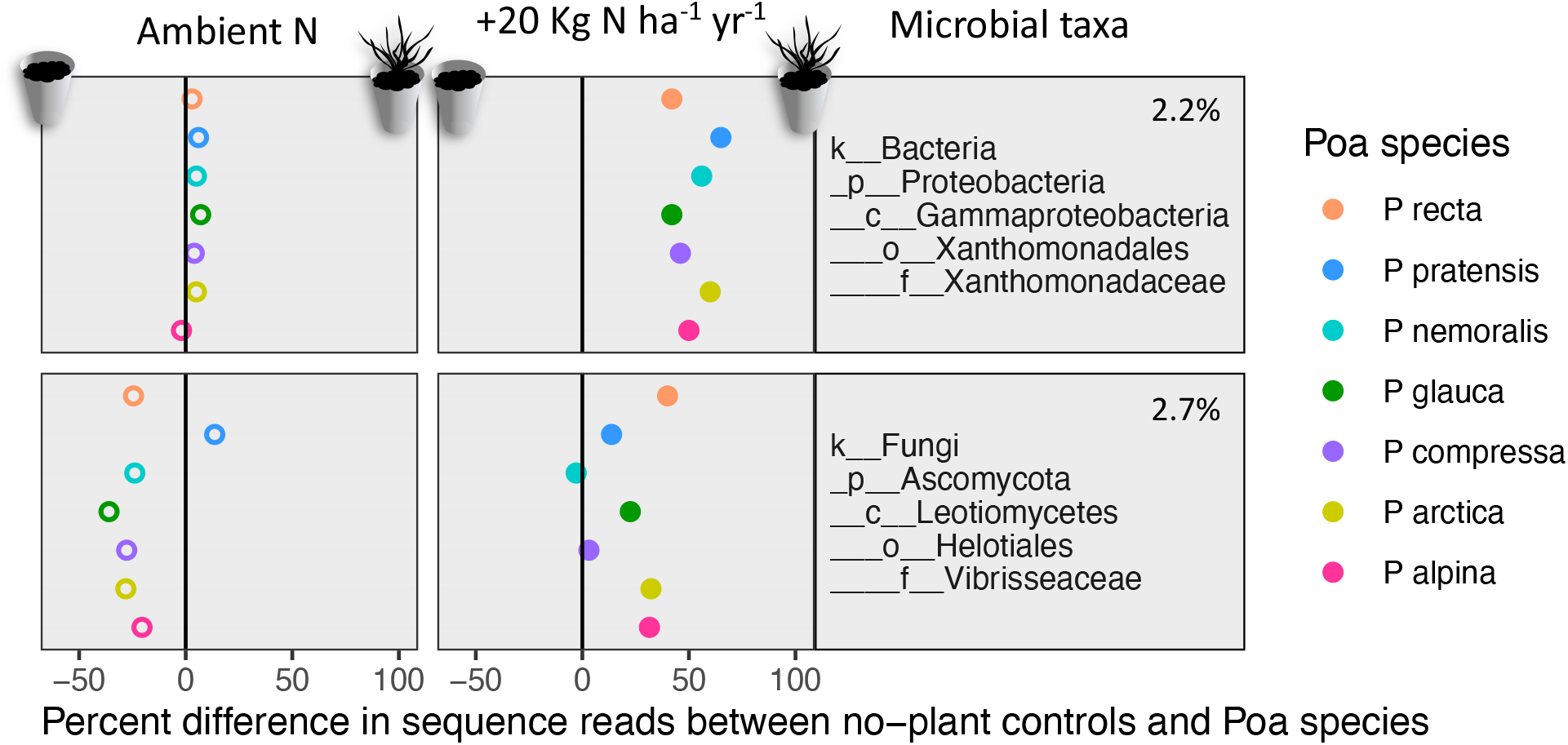
A fungal family (top) and a bacteria family (bottom) where points represent the mean relative abundance in pots with a particular *Poa* species after subtracting the mean relative abundance of the family from the no-plant control pots. The column on the left shows whether families are more likely to be found in *Poa* rhizospheres or soil not influenced by roots whereas the column on the right shows how N addition influences the relative abundance of families. Phylum, class, order, and family are provided for each family, and the percent of reads from each family in the entire dataset are provided with the taxonomic name. See Figs. S2 and S3 for more microbial families.

To account for the potential effects of soil pH on microbial community composition, we evaluated how variance in community composition that could be explained by N versus pH. This test was conducted because soil pH was 0.5 units lower on average in N addition pots compared to ambient N pots. Variance partitioning revealed that 5% of the explained variance in microbial community composition could be attributed to N addition while only 1% of the variation could be attributed to variation in pH. Both N and pH together explained an additional 6% of the variation in microbial community composition. This affirmed our ability to interpret the direct effects of the N treatment on microbial communities.

After establishing that N had unique effects on *Poa* species (Fig. S5), we were able to test whether plant growth responsiveness to N among *Poa* species (difference in plant height between N treated plants and ambient N treated plants) was related to the magnitude of microbial community change in response to N addition (Bray Curtis dissimilarities between N treated communities and ambient N treated communities)(Fig. S2). Neither fungal nor bacteria communities could be explained by plant responses to N within nor among *Poa* species (Table 4, Fig. 3).

**Table 4.**
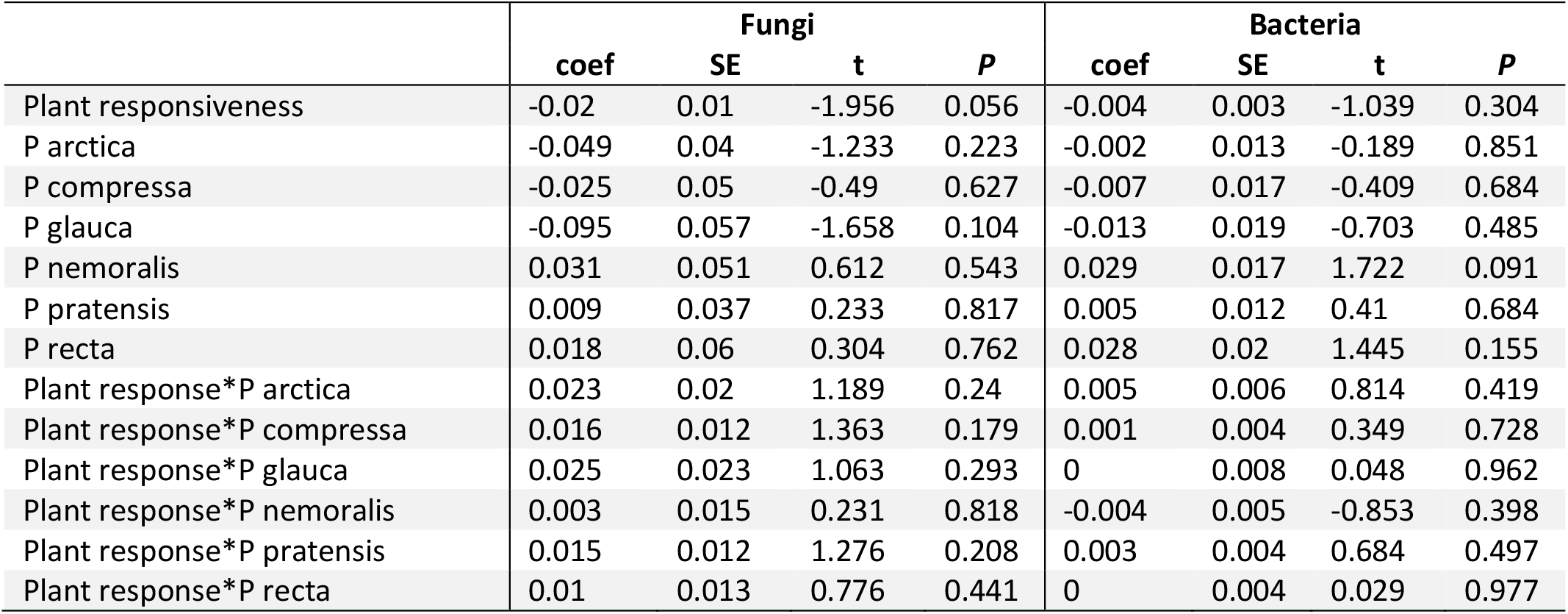
Linear regression results indicate that the magnitude of microbial community response to N addition is not explained by plant responses to N (change in plant height) while accounting for *Poa* species differences.

|  | Fungi |  |  |  | Bacteria |  |  |  |
| --- | --- | --- | --- | --- | --- | --- | --- | --- |
|  | coef | SE | t | P | coef | SE | t | P |
| Plant responsiveness | -0.02 | 0.01 | -1.956 | 0.056 | -0.004 | 0.003 | -1.039 | 0.304 |
| P arctica | -0.049 | 0.04 | -1.233 | 0.223 | -0.002 | 0.013 | -0.189 | 0.851 |
| P compressa | -0.025 | 0.05 | -0.49 | 0.627 | -0.007 | 0.017 | -0.409 | 0.684 |
| P glauca | -0.095 | 0.057 | -1.658 | 0.104 | -0.013 | 0.019 | -0.703 | 0.485 |
| P nemoralis | 0.031 | 0.051 | 0.612 | 0.543 | 0.029 | 0.017 | 1.722 | 0.091 |
| P pratensis | 0.009 | 0.037 | 0.233 | 0.817 | 0.005 | 0.012 | 0.41 | 0.684 |
| P recta | 0.018 | 0.06 | 0.304 | 0.762 | 0.028 | 0.02 | 1.445 | 0.155 |
| Plant response*P arctica | 0.023 | 0.02 | 1.189 | 0.24 | 0.005 | 0.006 | 0.814 | 0.419 |
| Plant response*P compressa | 0.016 | 0.012 | 1.363 | 0.179 | 0.001 | 0.004 | 0.349 | 0.728 |
| Plant response*P glauca | 0.025 | 0.023 | 1.063 | 0.293 | 0 | 0.008 | 0.048 | 0.962 |
| Plant response*P nemoralis | 0.003 | 0.015 | 0.231 | 0.818 | -0.004 | 0.005 | -0.853 | 0.398 |
| Plant response*P pratensis | 0.015 | 0.012 | 1.276 | 0.208 | 0.003 | 0.004 | 0.684 | 0.497 |
| Plant response*P recta | 0.01 | 0.013 | 0.776 | 0.441 | 0 | 0.004 | 0.029 | 0.977 |

**Fig. 3.**
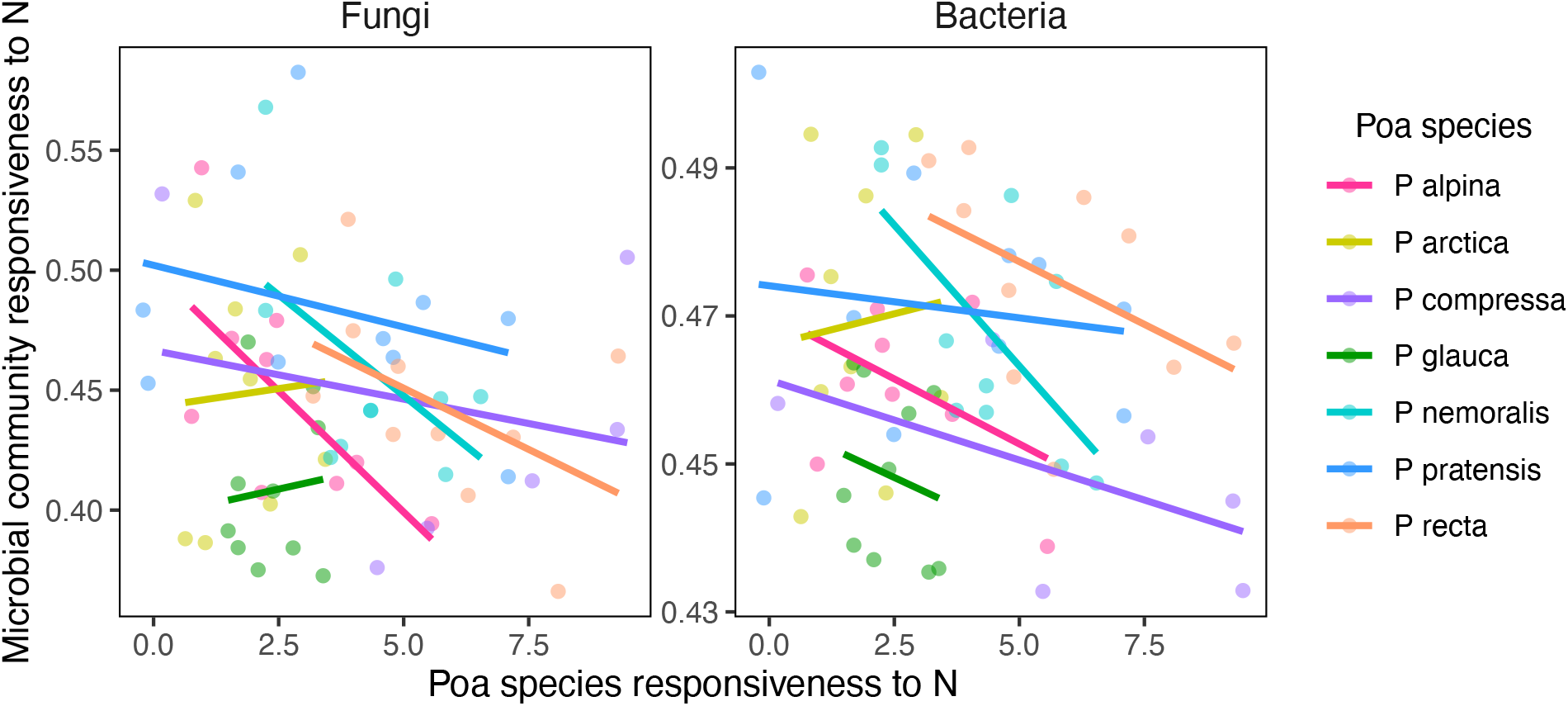
Relationship between plant growth responsiveness to N (difference in plant height between ambient N and N addition pots) and the magnitude of microbial community compositional response to N (Bray Curtis dissimilarities between ambient N and N addition pots).

## Discussion

Our objective in this study was to investigate how congeneric taxa influence soil microbial communities and whether species-specific plant effects depend on N availability which can vary substantially in natural systems due to atmospheric N deposition and in agricultural systems that vary in fertilization rates. We found that *Poa* species uniquely affected fungal but not bacterial rhizosphere communities, that N availability mediated the plant species effect on both fungal and bacterial communities.

### Plant species’ effects on microbial composition

First, we hypothesized that congener plant species have unique effects on microbial communities and that some of the variation in microbial communities among these congeners can be explained plant species’ traits and/or phylogeny, a proxy for unmeasured traits. As previously stated, fungal communities differed among *Poa* species while bacterial communities did not under ambient N. Thus, we conclude that congeneric *Poa* species can indeed have unique, non-random effects on microbial communities. The effect is consistent among replicates (small *P*-value) but the strength of the effect is relatively small (low R^2^) and influenced by changes in the relative abundance of rare taxa, as has been shown elsewhere (Dawson et al. 2017).

Differences in *Poa* species’ influence on microbial composition were unrelated to plant trait variation among *Poa* species and phylogeny (a proxy for unmeasured plant traits). We offer three potential reasons why our chosen predictors were poor indicators of plant effects on soil microbes. First, the *Poa* species we used may be too physiologically similar to detect differences in their effects on soil microbial communities. We examined non-sister taxa in the same genus to capture an intermediate level of similarity on the continuum of relatedness examined in previous studies (Bardgett et al. 1999; Zancarini et al. 2012; Bouffaud et al. 2014; Barberán et al. 2015). While our results do not provide support for genera (i.e. interspecific relationships) representing the necessary balance between evolutionary constraints and ecological divergences needed to detect phylogenetic signal in microbial community composition, it is possible that other plant genera may possess the “just right” amount of variation in how they interact with root-associated soil.

Second, phylogeny may not be a good proxy for the unmeasured plant traits that are relevant to rhizosphere interactions (Reinhart and Anacker 2014; Leff et al. 2018). While phylogenetic signal was found in plant-soil feedbacks in one study that robustly quantified feedbacks within multiple plant clades (Anacker and Strauss 2016), no phylogenetic signal was found in other studies that did not measure feedbacks (Mehrabi and Tuck 2015; Fitzpatrick et al. 2017). We considered that the feedback effect may involve relatively few microbial taxa, in which case our investigation of entire microbial communities could have obscured the signature of *Poa* species associations with specific microbial taxa. We also found that the *Poa* phylogeny was also not a good proxy for our *measured* plant traits (i.e, phylogenetic signal tests all P > 0.05; not shown), reflecting either a lack of trait conservatism or error in the phylogeny estimation or trait measurement. Taken together, these results indicate that *Poa* phylogenetic relatedness may be a poor predictor of plant traits and plant effects on soil microbial communities.

Third, neither plant traits (plant height, dry shoot mass, dry root mass, and SLA) nor the plant species range-wide traits (soil pH and mean annual temperature in *Poa* species’ geographical ranges) appear to be useful indicators for the unmeasured factors (e.g. exudate chemical composition (Haichar et al. 2014)) that may have driven the observed divergence in microbial community composition among *Poa* species (Cantarel et al. 2014; Moreau et al. 2015). While the traits we chose are associated with plant resource use and tradeoffs in life history strategies (Funk et al. 2017), they do not appear to explain differences in plant species’ allocation of resources to exudate composition or quantity. A different pot experiment also found above-ground plant traits and plant phylogeny to be poor indicators of plant effects on soil microbes for diverse grassland plant species (Leff et al. 2018). A third study on distantly related grassland species also found no phylogenetic signal in plant effects on soil microbes but did find that SLA was a significant predictor of plant-soil feedback (Fitzpatrick et al. 2017). Together, commonly studied above-ground plant traits usually lack the capacity to detect differences in below-ground ecology among plant species.

### Nitrogen addition influences Poa species effects on soil microbial communities

We found that N addition enhanced species-specific plant effects on both fungal and bacterial community composition. Nitrogen addition has previously been shown to moderate plant effects on bacteria community composition when tested on distantly related plant species (Frank and Groffman 2009; Moreau et al. 2015). In our study, bacterial community composition was significantly different among *Poa* species only under elevated N but not in ambient N pots. This finding suggests that even when N is abundant plant-microbe interactions may be important for plant or microbial nutrient acquisition as opposed to plants and microbes operating more independently when neither N nor C are scarce (Kaye and Hart 1997; Bell et al. 2015). We also found that N addition had a stronger direct effect on bacterial and fungal community composition than indirect effects of decreasing pH associated with the N addition. This result has also been described in other studies (Frederick and Klein 1994; Groffman et al. 1996; Innes et al. 2004; Farrer et al. 2013). Many factors influence how much N addition influences pH and the associated microbial response to pH versus N including the natural acidity of the soil and the rate of N applied (Stevens et al. 2004; Geisseler and Scow 2014).

To address whether plant species’ growth responses to N addition (or all plants across species) influenced the magnitude of change in microbial community composition in response to N, we tested for a positive linear relationship between plant growth (difference in plant height between N treated plants and ambient N treated plants) and microbial community composition change (Bray Curtis dissimilarities between N treated communities and ambient N treated communities). Our results did not support the hypothesized relationship between plant growth responses and the magnitude of change in microbial community composition. Plant growth was hypothesized to correlate with belowground plant effects on soil microbial composition because plant growth responses to N not only vary among species (Wardle et al. 2004; Rinnan et al. 2007) but plants also alter allocation of resources to growth and exudates when N availably changes (Bowsher et al. 2017). If the change in types and quantities of C molecules is large enough, microbial populations will grow or shrink in response to resources and ultimately alter community composition (Paterson et al. 2007). In our study, *Poa* species’ allocation of C for plant growth may have resulted in less root exudation or little change from ambient N conditions. Therefore, fast-growing plant species in the genus *Poa* may not necessarily invest additional C belowground to stimulate microbial acquisition of limiting resources for sustained growth. While we examined taxonomic composition to quantify the microbial community response, Rinnan 2007 found that greater microbial biomass was positively correlated with plant biomass (but see Bardgett et al. 1999). N availability has diverse effects on rhizodeposition among studies, even after accounting for variation in C pools and measurement units across studies (Bowsher et al. 2017). Indeed, while many factors influence rhizodeposition, our experimental manipulation—which isolated plant species and N effects—provides context for ongoing and future research that aims to identify mechanisms by which plant species affect soils differently, including differentiating key elements of plant exudate profiles or the genes that regulate exudate production.

## Conclusion

Our study concludes that plant congeners can have distinct effects on rhizosphere microbial composition, although basic plant traits do not appear to be useful indicators. Our finding that N increase accentuates species-specific effects reveals important context dependency in plant-microbe interactions. This context dependency suggests that previous interpretation of functional relationships between plants and microbes in the rhizosphere may only be valid for the specific N availability of a study. If additional plant clades exhibit similar N dependent associations, this knowledge could be used to more accurately forecast plant responses and perhaps ecosystem responses to variation in N supply.

## Declarations

### Availability of data and material

Data will be submitted to the Long Term Ecological Research (LTER) public repository and sequences will be submitted to NCBI

### Code availability

Analyses were conducted using open access R packages and no novel coding approaches were used.

## Acknowledgments

We thank Noah Fierer, Katheryn Suding, Nichole Barger, Eve-Lynn Hinkley and Jonathan Leff for insightful input throughout the study; Jessica Henley, Thomas Lemieux, Janice Harvey, Tim Hogan, Dina Clark, Ben Murphy, Matt Gebert, Holly Archer, Max Owens, Tyler Justice for help with the experiment and/or data collection; Tobin Hammer, Josh Grinath, and Richard Lankau for statistical input. We thank Black Dog LED, Boulder, CO for donation of an LED light panel for this experiment, and the EBIO department’s graduate student Writing Co-op, and QDT for writing and analysis support. This research was funded by Beverly Sears Fund, Jon Marr Memorial Fund, the Undergraduate Research Opportunity Program, and the EBIO Research Grant at the University Colorado.

## Supplementary Information

**Table S1.** Species by gene matrix with genes used to calculate phylogenetic distance among *Poa* species

| Taxon | ets | its | matk | rbcl | trnT-trnL |
| --- | --- | --- | --- | --- | --- |
| *Poa alpina* | 297375143 | 297375339 | 459929648 | 459994671 | 86169391 |
| *Poa arctica* | 297375147 | 297375343 | 371533758 | 459994755 | 86169414 |
| *Poa compressa* | 685211931 | 209887699 | 744393645 | 685212046 | 86169408 |
| *Poa glauca* | 297375180 | 209887701 | 459929766 | 459994799 | 86169409 |
| *Poa nemoralis* | 297375211 | 297375385 | 607345128 | 607344860 | 297375296 |
| *Poa pratensis* | 297375225 | 297375398 | 459929844 | 459994878 | 86169416 |
| *Poa reflexa* | NA | 297375399 | NA | NA | 297375306 |

**Fig. S1.**
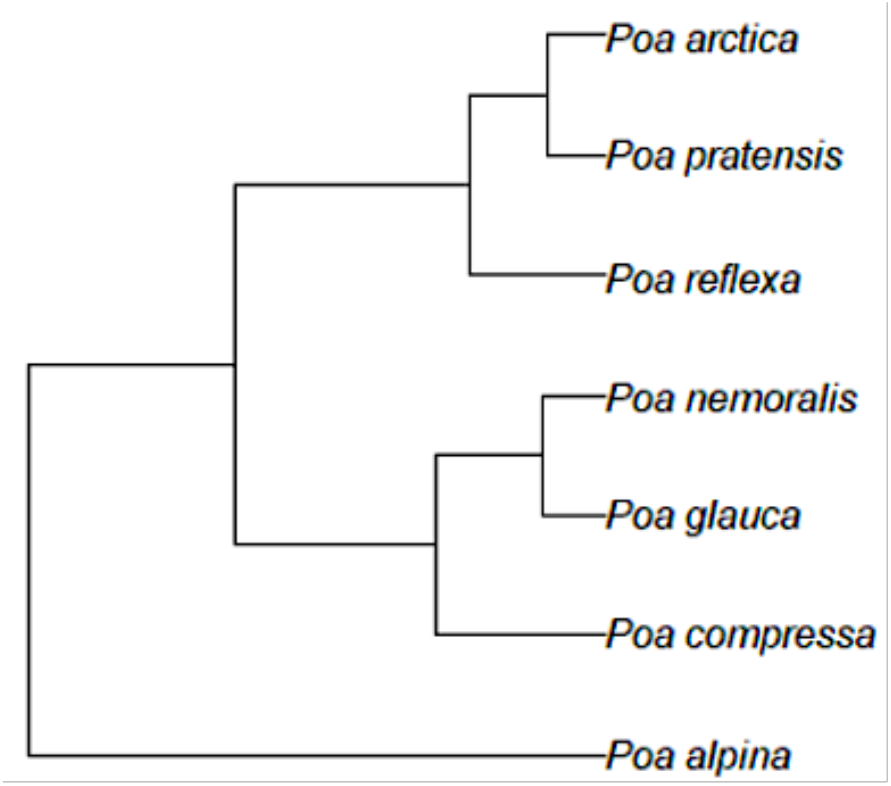
Pruned *Poa* phylogeny showing evolutionary relationships among the seven *Poa* species in the experiment. The phylogeny is based on the topology from Gillespie et al. 2007 with branch length calibration based on the gene regions shown in Table S1.

**Table S2.**
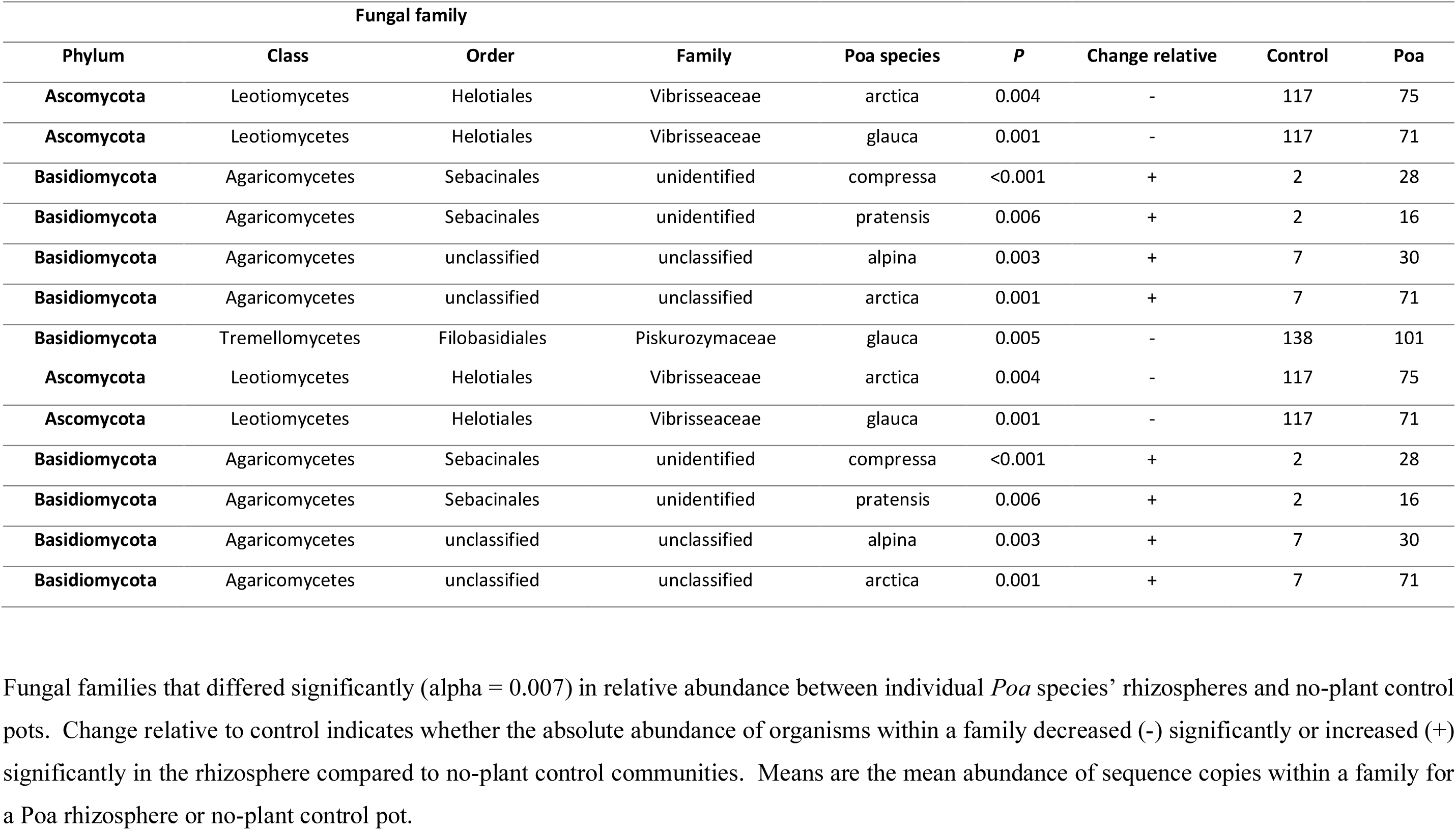
Fungal families affected by Poa species compared to no-plant controls.

**Table S3.**
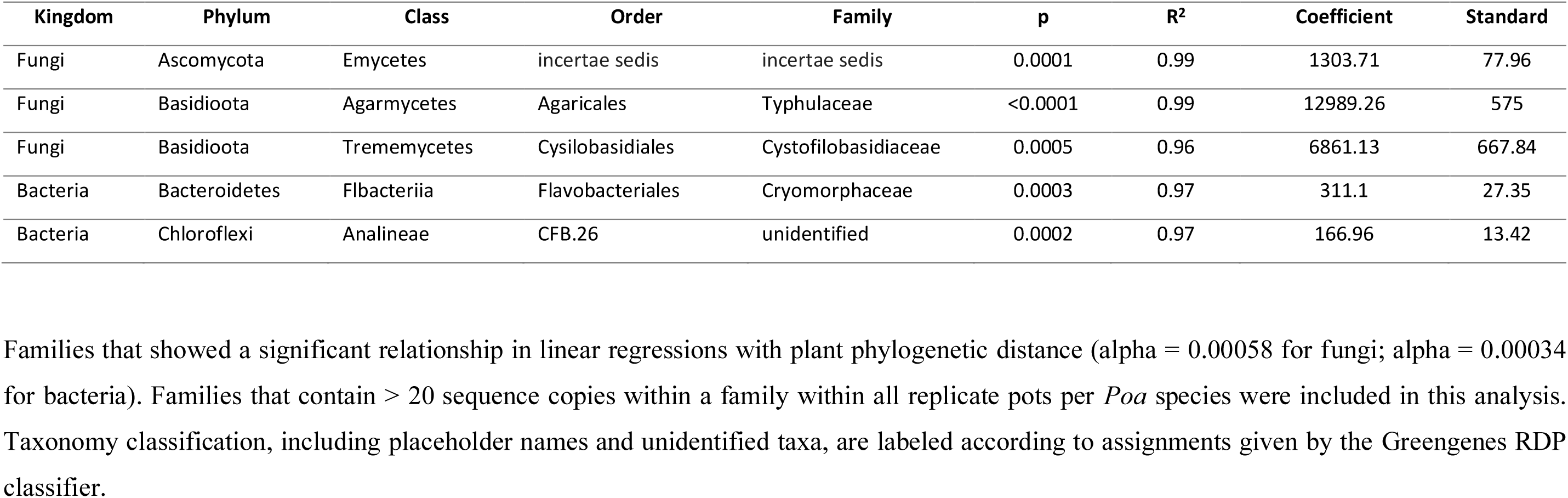
Fungal and bacterial families with a significant relationships with phylogenetic distance among Poa species.

**Fig. S2.**
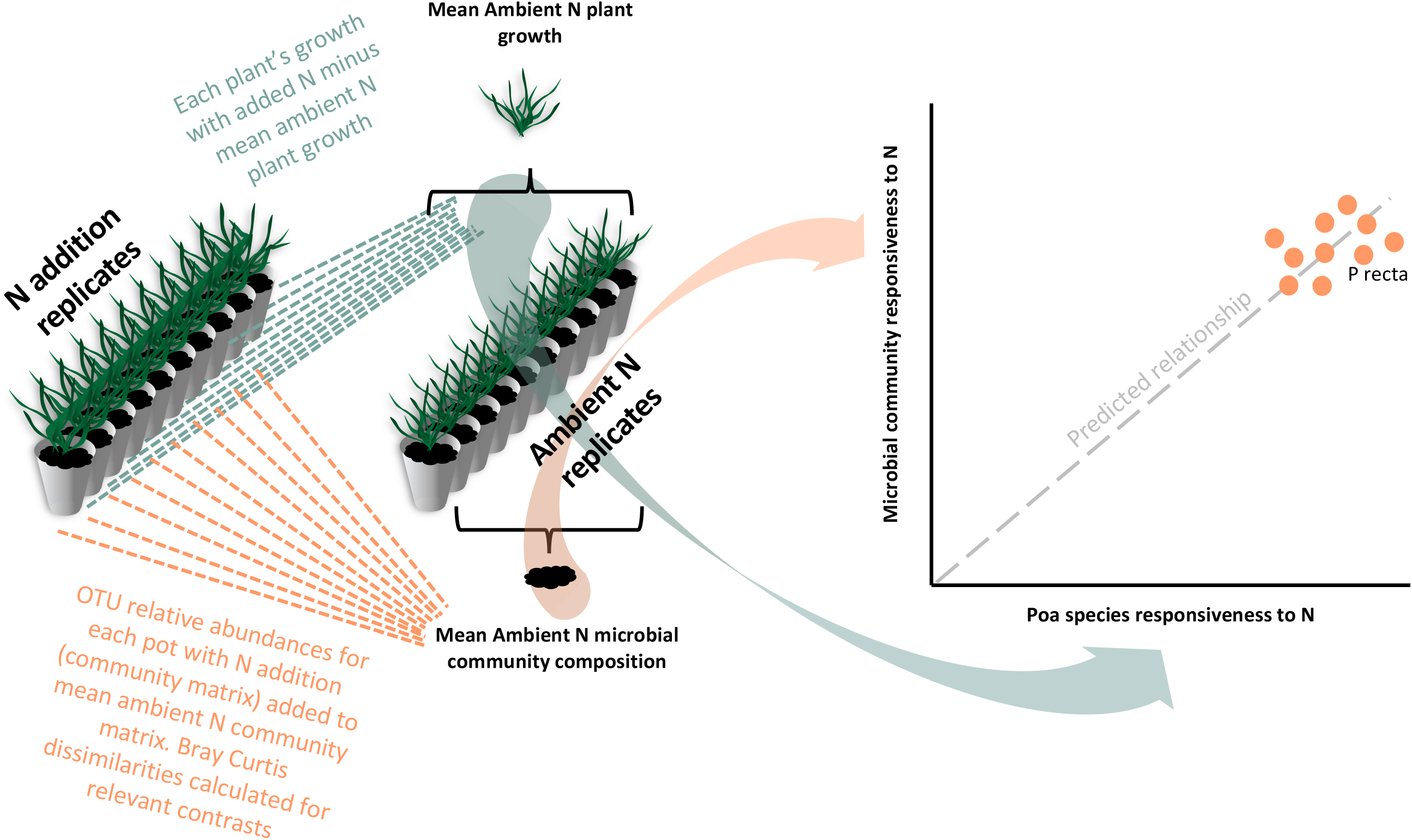
Example calculation of plant and microbial community responsiveness to N with *P recta* to test final hypothesis that there is a relationship between plant and microbial responses to N addition.

**Fig. S3.**
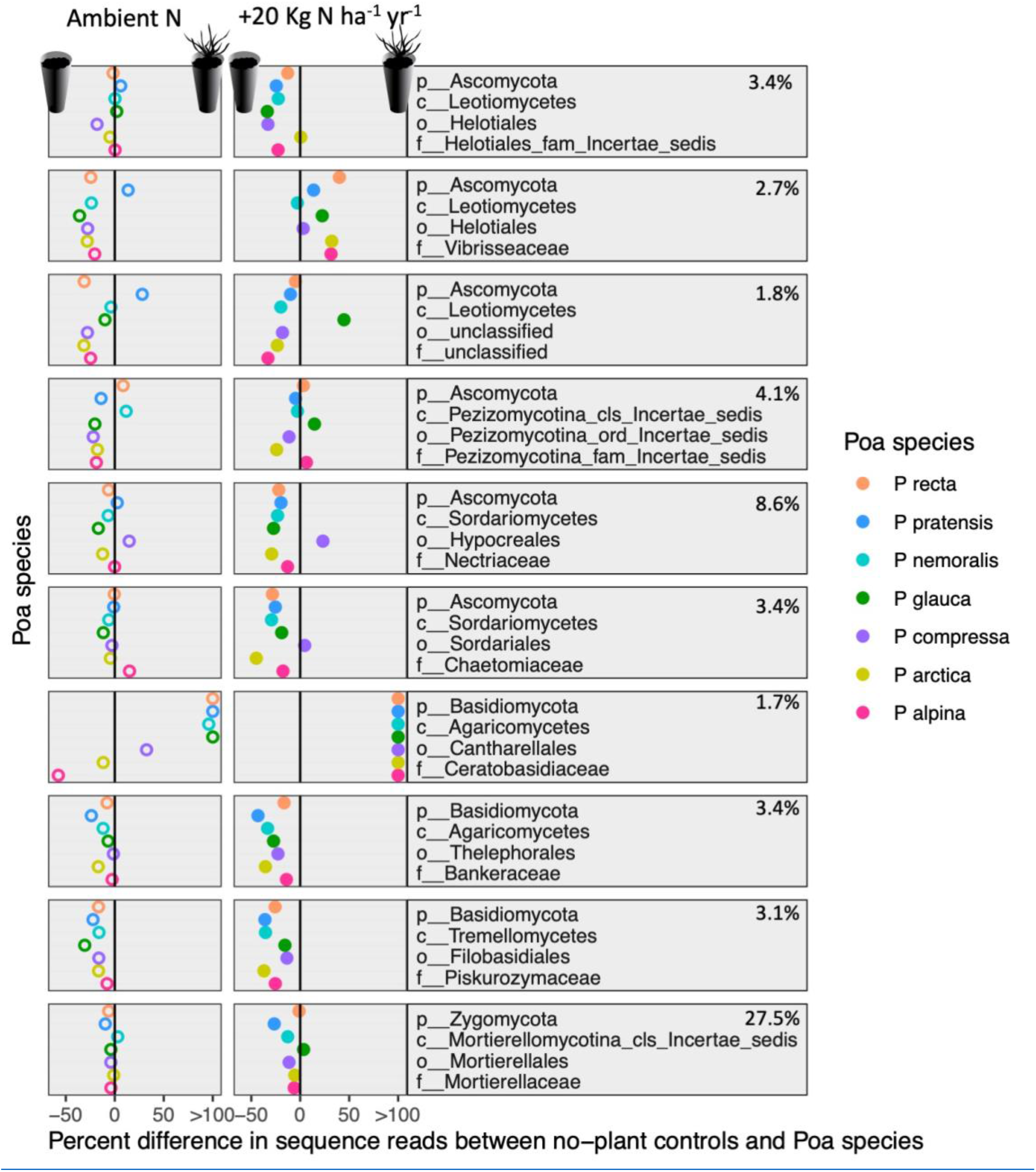
Ten fungal families with highest total relative abundance in the greenhouse experiment are shown. Points represent the mean relative abundance in pots with a particular Poa species after subtracting the mean relative abundance of the bacterial family from the no-plant control pots. The column on the left shows whether families are more likely to be found in Poa rhizospheres or soil not influenced by roots whereas the column on the right shows how N addition influences the relative abundance of families. Phylum, class, order, and family are provided for each family, and the percent of reads from each family in the entire dataset are provided with the taxonomic name.

**Fig. S4.**
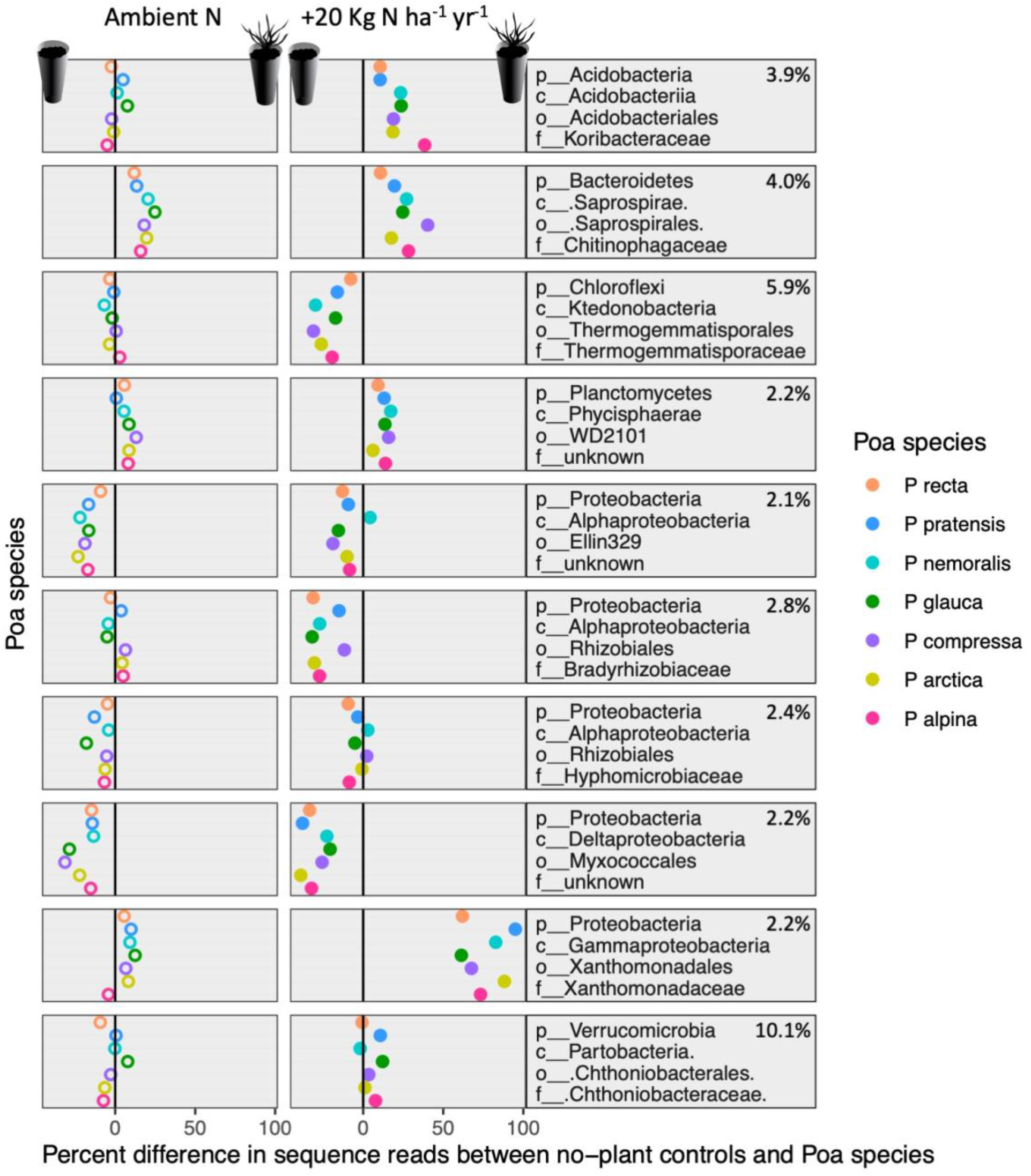
Ten bacterial families with highest total relative abundance in the greenhouse experiment are shown. Points represent the mean relative abundance in pots with a particular Poa species after subtracting the mean relative abundance of the bacterial family from the no-plant control pots. The column on the left shows whether families are more likely to be found in Poa rhizospheres or soil not influenced by roots whereas the column on the right shows how N addition influences the relative abundance of families. Phylum, class, order, and family are provided for each family, and the percent of reads from each family in the entire dataset are provided with the taxonomic name.

**Fig. S5.**
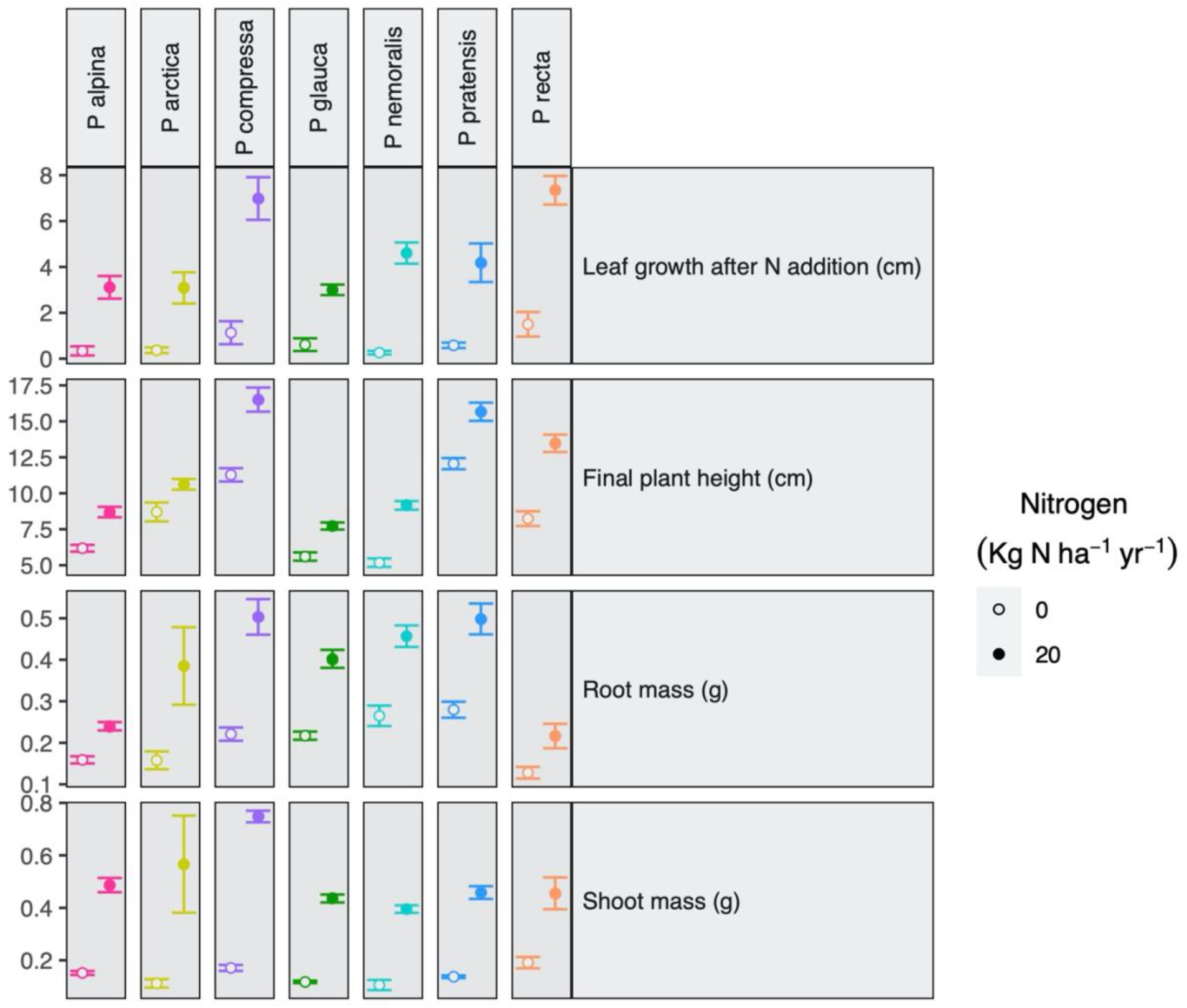
Effect of N on plant measurements shown as means and standard error of the means for ambient and N addition pots. All contrasts between N addition and ambient N replicates are statistically significant for each Poa species with *P* < 0.04 for all tests.

**Fig. S6.**
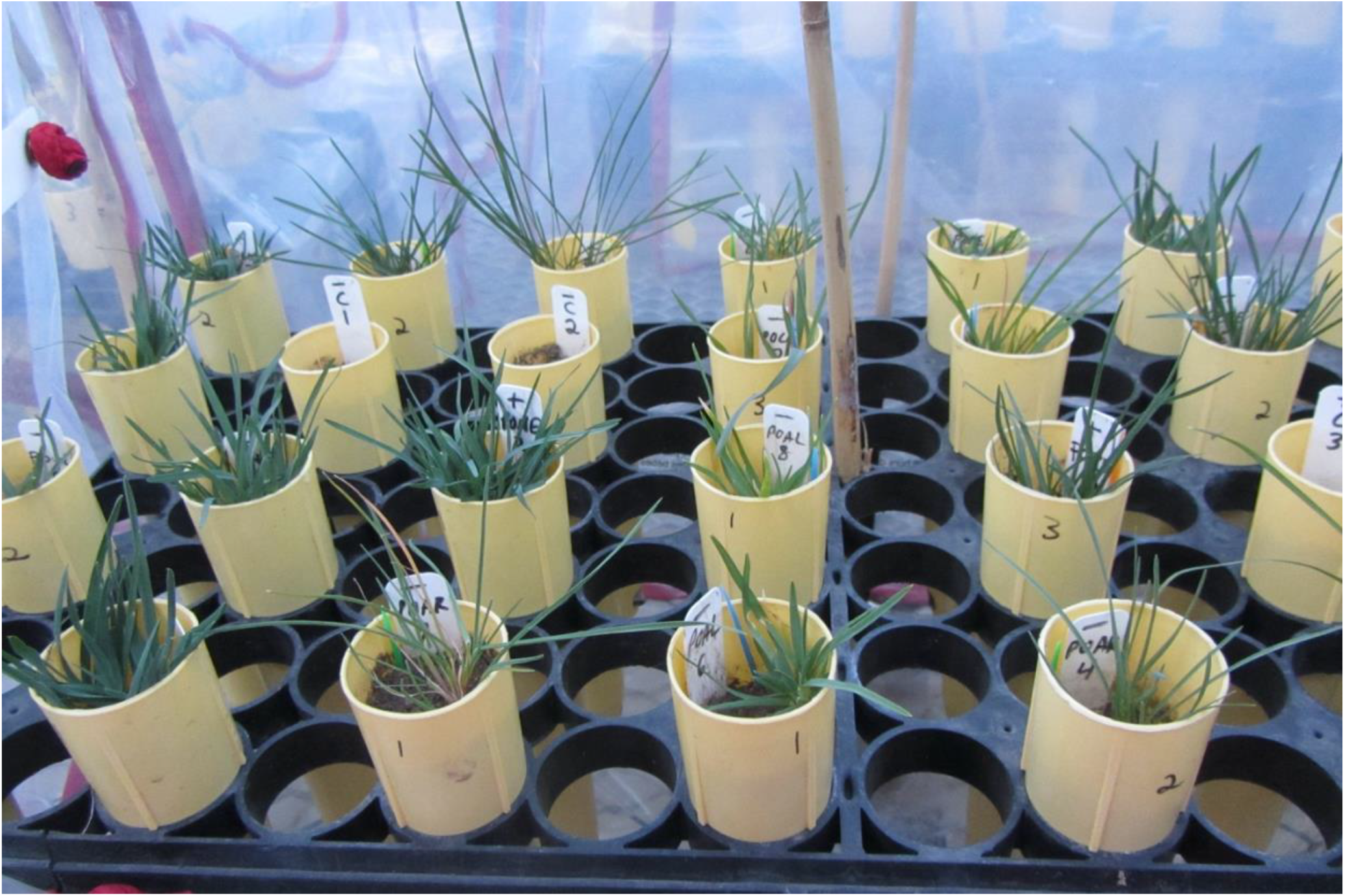
Image of *Poa* grasses growing in cone-tainers in greenhouse experiment.

